# Stretching of long double-stranded DNA and RNA described by the same model

**DOI:** 10.1101/2022.03.09.483706

**Authors:** Alexander Y. Afanasyev, Alexey V. Onufriev

## Abstract

We propose a bead-spring model that accurately reproduces a variety of experimental force-extension curves of long double-stranded DNA and RNA, including torsionally constrained and unconstrained DNA, and negatively supercoiled DNA. A key feature of the model is a specific non-convex energy function of the spring. We provide an algorithm for obtaining five required parameters of the model from experimental force-extension curves. In the plateau region of the force-extension curves, our molecular dynamics simulations show that the polymer separates into a mix of weakly and strongly stretched states without forming macroscopically distinct phases.

## 1 Introduction

Double-stranded (ds) DNA and RNA molecules are constantly under different loading conditions in living cells.^1–3^ Knowledge of the mechanical properties of these polymers is of great importance for understanding many key cellular processes. ^4–7^ A widely used experimental technique employed to explore the mechanics of polymers is the single-molecule stretching with optical or magnetic tweezers.^5,8–10^

To interpret these experiments under low tension, several variants of the worm-like chain model (WLC) have gained popularity.^8,11–16^ The popularity of the models stems, in large part, from their conceptual simplicity, the clear physics behind them, the direct connection with the key mechanical parameters of the polymers, and the facility for extracting these properties from the experimental data. For example, according to the extensible worm-like chain (eWLC) model,^8,17,18^ at a given stretching force F, the extension *L*_*x*_ = *L*_0_(1 + *F/S* − (*k*_*B*_*T/*4*FP*)^1*/*2^), where *k*_*B*_ is the Boltzmann constant, and T is the absolute temperature. Thus, the parameters of the polymer, such as the polymer persistence length P, the stretch modulus S, and its contour length *L*_0_ (its maximum end-to-end distance), can be obtained by fitting the model to the experimental force-extension curve. ^8,19^

However, when it comes to dsDNA and dsRNA under high tension, these popular models have a serious limitation. Specifically, the single-molecule experiments reveal a peculiar plateau region on force-extension curves for dsDNA^8,20–22^ and dsRNA.^8,23^ Only the stretching behavior of the molecules below the plateau is well described by the eWLC model,^8,17,18^ but the plateau itself remains out of reach of this otherwise appealing model (see *e*.*g*. Figure 5 of the “Results” section).

To describe full force-extension curves of dsDNA, including the plateau, a variety of theoretical models have been proposed.^24–27^ These models are based on statistical mechanics, and describe a number of possible states with parameters fitted to experimental data. Despite the clear advantage of being more general than the variants of the WLC, these more complex models have not been used nearly as widely in interpreting single-molecule experiments, possibly because the connection to macroscopic parameters that describe properties of the polymers is no longer as intuitive as in the variants of the WLC.

On the other hand, molecular dynamics models, fully atomistic^28–40^ or coarse-grained (mesoscopic),^41–46^ provide different insight into the mechanics of the nucleic acids. The specific choice of the level of granularity of the models depends on the problem at hand. In general, these models rely on empirical potential functions (force-fields) that describe interactions between atoms or beads through a combination of bond stretching, bending, torsional, and electrostatic potentials, to name a few, where each potential has multiple fitted parameters, making the total number of parameters as many a hundred for some general-purpose biomolecular force-fields.^47,48^ An advantage of these models, compared to fully phenomenological models such as the variants of the WLC, is that they can provide a microscopic, including atomic-level, interpretation of experiment. The drawback is that direct connection to the macroscopic parameters that describe polymer deformation is harder to establish, and the simulation outcomes may depend on the force-field chosen. These complex force-fields are fine-tuned to work with all of their components present, making it difficult to ask questions that require turning various physical interactions on or off. For the specific case of the DNA and RNA, several of these models have enough complexity built into them to reproduce, to various degrees, the overstretching plateau region, which is a clear advantage over the variants of the WLC. A number of simulation studies of dsDNA and dsRNA stretching at this level of description has been performed,^49–54^ but still we are unaware of a mesoscopic or a fully atomistic model that accurately reproduces various regions of experimental force-extension curves of both long dsDNA and dsRNA within the same model.

We suggest a mesoscopic model that may be considered a hybrid between the two approaches, with just enough complexity to faithfully account for the plateau region and its basic physics. We aim to retain the appealing simplicity level of the variants of the WLC with its direct connection to mechanical properties of the polymer, while also retaining the most basic feature of the molecular simulation approach – microscopic granularity. Our longterm motivation is to construct a model that may potentially replace or at least complement the variants of the WLC in the routine interpretation of single-molecule experiments.

## 2 Results and discussion

We propose a mesoscopic model, based on a non-convex potential energy function, that reproduces, closely, the experimental stretching behavior of long dsDNA and dsRNA with one plateau region on their force-extension curves. We use the model to explore the possibility of a phase separation in the plateau region.

The conceptual basis of the proposed model is a recently proposed general framework,^49,55^ in which the existence of the plateau region on the force-deformation curve stems from a non-convex profile of the energy of the system. Namely, if the energy as a function of deformation is not convex over some interval, then the system can minimize its energy by separating into two states: weakly and strongly deformed, with the geometric characteristics of the states corresponding to the beginning and the end of the convex hull of the energy function, respectively. ^49,55^

Our proposed model treats the dsDNA or dsRNA molecule as a chain of *N* beads connected by non-linear springs, Figure 1A. The stretching behavior of the non-linear spring is governed by the potential, *U*(*r*), in the form of a piecewise continuously differentiable function:

**Figure 1:**
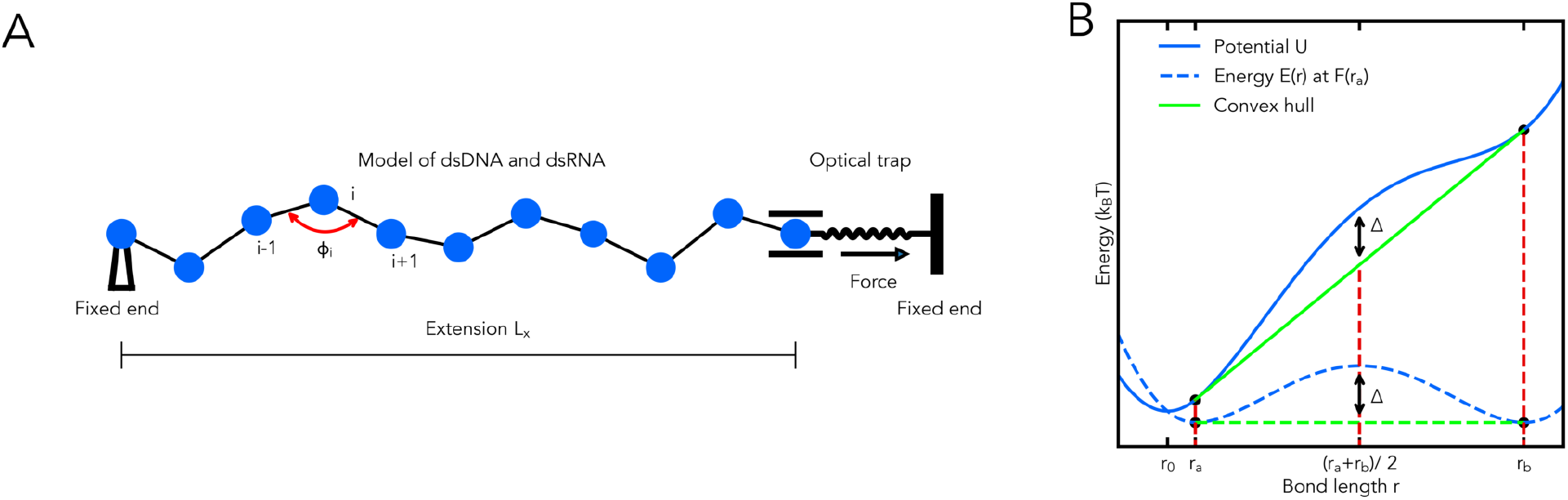
**(A)** A schematic showing the bead-spring model, a model of an optical trap and applied displacement boundary conditions on the ends of the chain. The leftmost bead of the chain is fixed, and the rightmost bead is allowed to move in the horizontal direction only. **(B)** The non-convex potential energy *U* as a function of *r*. The existence of a non-convex region in *U*(*r*), controlled by the parameter Δ, leads to non-uniform stretching of the chain. The potential energy of the spring at an applied load along the *r*-direction of *F*(*r*_*a*_) = *U′*(*r*_*a*_), *E*(*r*) = *U*(*r*) − *F*(*r*_*a*_)*r*, is shown as the dashed blue curve.

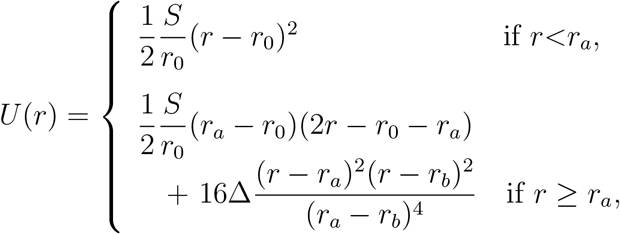

where *S* is the stretch modulus of the molecule (in *k*_*B*_*T/Å*); *r*, the distance between the centers of neighboring beads - the bond length (in Å); *r*_0_, the equilibrium bond length (in Å); *r*_*a*_, the bond length at the beginning of the convex hull (in Å); *r*_*b*_, the bond length at the end of the convex hull (in Å); Δ, the energy difference between the potential *U* (in *k*_*B*_*T*) and the convex hull at *r*=(*r*_*a*_+*r*_*b*_)/2. Equivalently, Δ can be interpreted as the potential energy barrier height between two equally likely states of the spring – weekly and strongly stretched – at an applied load along the *r*-direction equal to the plateau force of *F*(*r*_*a*_) = *U ′*(*r*_*a*_), as depicted in Figure 1B.

To account for the bending of the chain, characterized by its persistence length *P*, we employ the discrete version of the WLC model, see “Methods”.

We propose an algorithm for extracting parameters of the new model, *P, S, r*_*a*_, *r*_*b*_, and Δ, from an experimental force-extension curve. The algorithm is based on approximating the force-extension curve by a piece-wise function 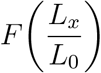, below. This function, the blue curve in Figure 2, describes the minimum energy state of the polymer chain with its fixed ends, governed by the stretching potential *U*(*r*).

**Figure 2:**
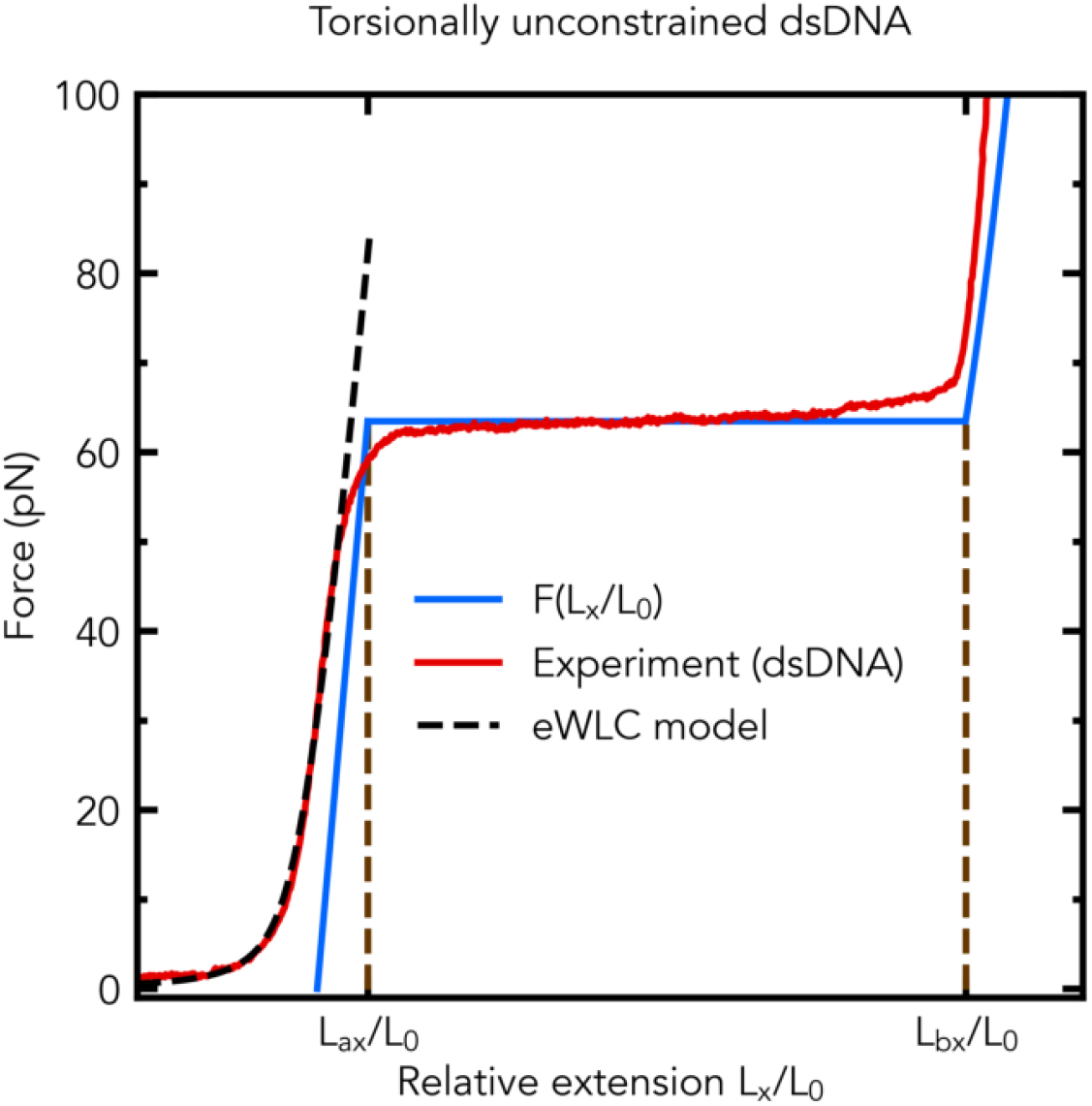
To derive parameters of the model (*P, S, r*_*a*_, *r*_*b*_, and Δ) from the experimental force-extension curve (red), we approximate the curve as an overlapping combination of two regimes: the eWLC (dashed black), which yields *S* and *P*, and one that follows the force-extension curve of the chain at the minimum energy state (blue), *F*(*L*_*x*_*/L*_0_), to obtain *r*_*a*_, *r*_*b*_ and Δ. The algorithm is illustrated here for torsionally unconstrained dsDNA (data are from Figure 5A of Ref. 19): the blue curve is *F*(*L*_*x*_*/L*_0_) with Δ = 8 *k*_*B*_*T, r*_0_ = 37.4 Å, *L*_*ax*_*/L*_0_ = 1.056, *L*_*bx*_*/L*_0_ = 1.72, *S* = 1133 pN, and the dashed black curve is the eWLC with *S* = 1133 pN and *P* = 471 Å.

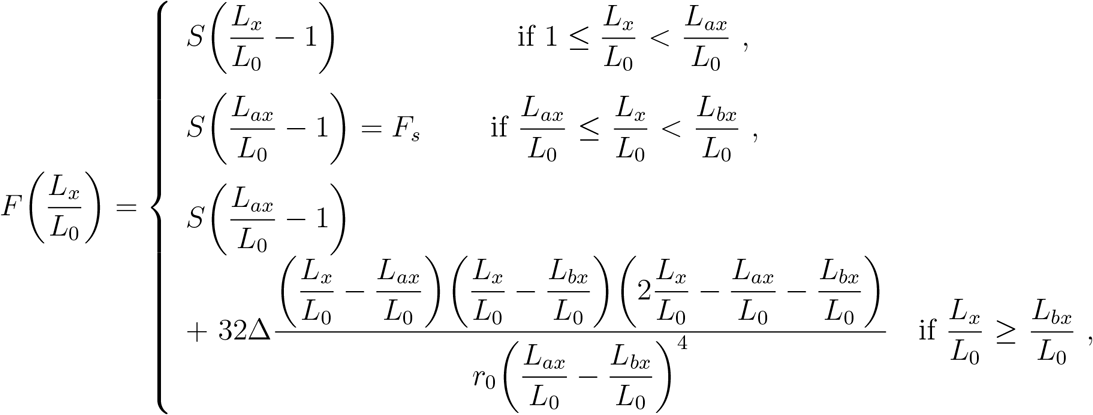

where *F*_*s*_ is the overstretching force at the middle of the plateau (in pN), *S* is the stretch modulus of the molecule (in pN), the energy difference Δ (in pN · Å), and the equilibrium bond length *r*_0_ (in Å). The approximation is reasonable for *L*_*x*_*/L*_0_ ≥ 1 when the chain behavior is not dominated by entropy effects.

The steps of the parameter extraction algorithm are as follows. First, extract *P* (in Å), *S* (in pN) from a fit of the eWLC model, *L*_*x*_*/L*_0_ = (1 + *F/S* − (*k*_*B*_*T/*4*FP*)^1*/*2^), to the experimental force-extension curve; note that the contour length of the molecule *L*_0_ and the equilibrium bond length *r*_0_ (in Å; potential *U*) are assumed known. Second, read *F*_*s*_, then, calculate *L*_*ax*_*/L*_0_ = 1 + *F*_*s*_*/S* and *L*_*bx*_*/L*_0_ = the end of the plateau + 0.02. Third, set *r*_*a*_ = *r*_0_*L*_*ax*_*/L*_0_ and *r*_*b*_ = *r*_0_*L*_*bx*_*/L*_0_. Finally, since *k*_*B*_*T* ≈ 41.41 pN · Å at *T* = 300 K, find Δ (in pN·Å) such that *F*(*L*_*x*_*/L*_0_) is parallel to the experimental curve for *L*_*x*_*/L*_0_ ≥ *L*_*bx*_*/L*_0_ (e.g., the blue curve in Figure 2).

We start exploring the model using coarse-grained Molecular Dynamics (MD) simulations, see “Methods”, by considering the case of torsionally unconstrained dsDNA in detail, Figure 3. The governing stretching potential *U*(*r*) is shown in Figure 3A for two different values of the parameter Δ; the other parameters of the model are obtained according to the algorithm. As illustrated in Figure 3B, the model clearly reproduces all the key features of the experimental force-extension diagram. The parameter Δ affects the transition of the chain to the plateau region, where the stretching of the chain is bi-modal, Figure 3B. The effect of larger Δ is a sharper transition to the plateau region in the force-extension diagram, as well as sharper peaks in the probability density *ρ* of relative bond lengths *r/r*_0_, as shown in Figure 3B.

**Figure 3:**
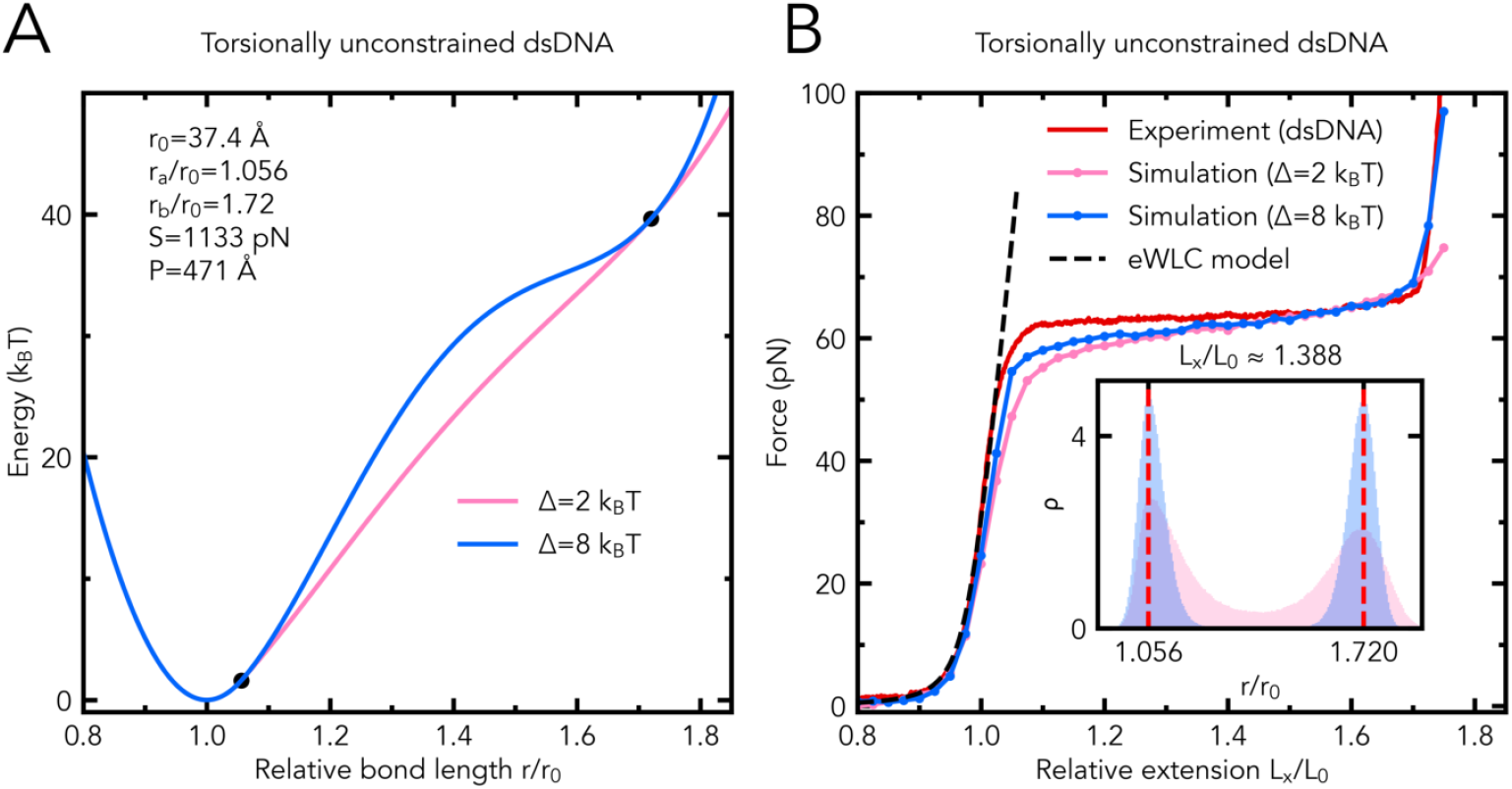
**(A)** The potential *U*(*r/r*_0_) of torsionally unconstrained dsDNA with Δ = 2 *k*_*B*_*T* (pink) and Δ = 8 *k*_*B*_*T* (blue). The parameters of *U*_*total*_ used in the simulations, are shown in the figure. **(B)** The results of simulations using the proposed model with Δ = 2 *k*_*B*_*T* (pink) and Δ = 8 *k*_*B*_*T* (blue) are compared with the experimental force-extention curve (data are from Figure 5A of Ref. 19) of torsionally unconstrained dsDNA and the eWLC model (parameters are in Figure 3A; the eWLC fit in the force range of 2-50 pN). Inset: the probability density *ρ* of relative bond lengths *r/r*_0_ for the model of torsionally unconstrained dsDNA with Δ = 2 *k*_*B*_*T* (pink) and Δ = 8 *k*_*B*_*T* (blue); *L*_*x*_*/L*_0_ ≈ 1.388.

In the plateau region, the polymer chain is a mix of two states – strongly and weekly stretched, Figure 4.

**Figure 4:**
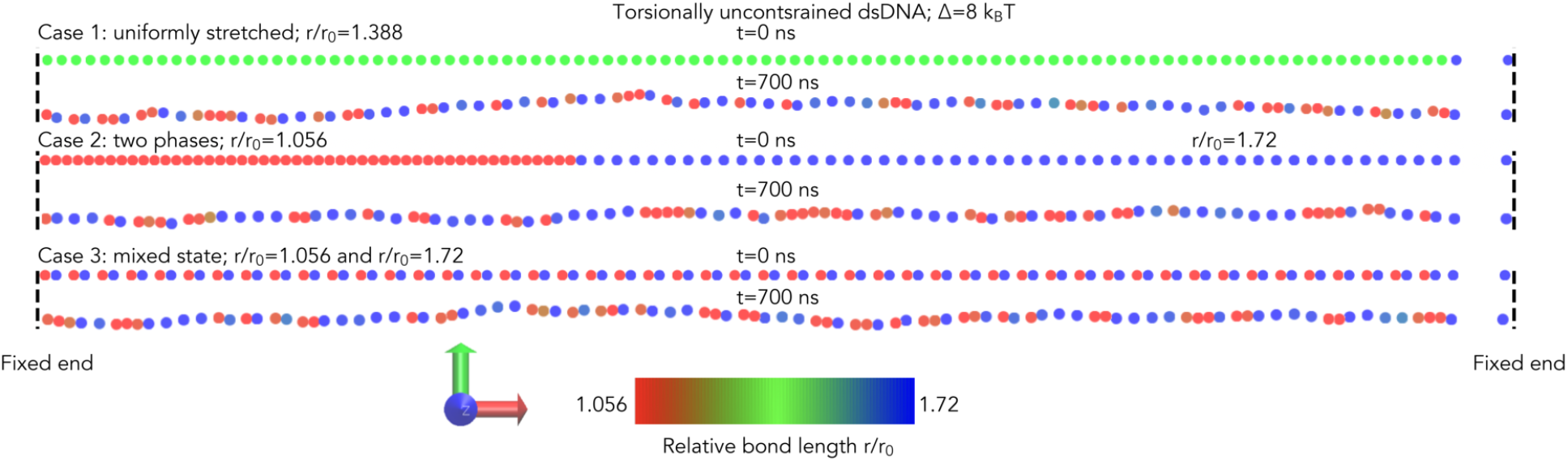
Conformations of the chain at t=0 ns and t=700 ns for torsionally unconstrained dsDNA with Δ = 8 *k*_*B*_*T*; *L*_*x*_*/L*_0_ ≈ 1.388. Case 1: the simulation begins from the ”one-phase” conformation, in which the beads are equally spaced with *r/r*_0_ = 1.388. Case 2: the simulation begins from the ”two-phase” conformation, in which the first 50 % of the beads are equally spaced with *r/r*_0_ = 1.056 and the other 50 % of the beads are equally spaced with *r/r*_0_ = 1.72. Case 3 (the Debye-Huckel potential is added): the simulation begins from the ”mixed state” conformation, in which the bond lengths alternate. The color scale (from red to blue) indicates the relative bond length between the bead of that color and its neighbor to the right. Snapshots are generated using VMD.

Further MD simulations based on the proposed model with the parameters obtained according to the algorithm, Figure 5, demonstrate that the model leads to a good agreement with the experimental force-extension curves of all tested dsDNA and dsRNA molecules, including, most importantly, the plateau region where the eWLC fails. In the plateau regions, the distributions (the probability density *ρ*) of relative bond lengths *r/r*_0_ demonstrate the co-existence of the two microscopic states: weakly and strongly stretched, Figure 5.

**Figure 5:**
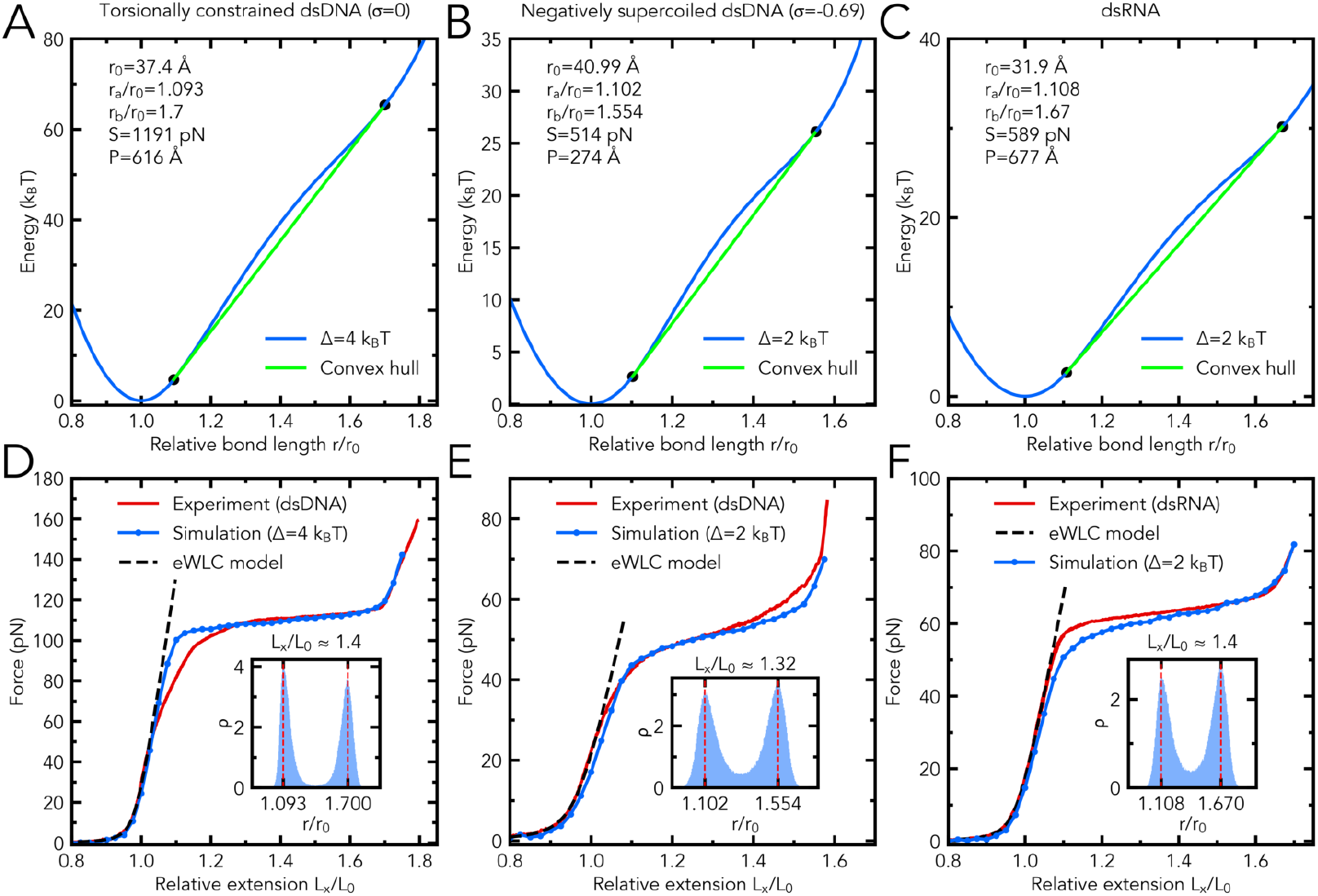
**(A, B, and C)**: The potential *U* as a function of *r/r*_0_ for **(A)** torsionally constrained dsDNA (superhelical density *σ* = 0), **(B)** negatively supercoiled dsDNA (*σ* = −0.69), and **(C)** dsRNA. The key parameters of *U*_*total*_ used in the respective simulations are shown in top left corner of each panel. **(D, E, and F)**: The results of simulations based on the proposed model are compared with the eWLC model (parameters are in Figure 5A, B and C) and the experimental force-extension curves of torsionally constrained dsDNA (*σ* = 0; the eWLC fit in the force range of 2-50 pN; experimental data are from Figure 5A of Ref. 19, negatively supercoiled dsDNA (*σ* = −0.69; the eWLC model, *L*_*x*_*/L*_0_ = *k*(1+*F/S* −(*k*_*B*_*T/*4*FP*)^1*/*2^), fits the experiment in the force range of 2-30 pN with *k*=1.096; experimental data are from Figure 5A of Ref. 19 with the values along the relative extension axis divided by *k*), and dsRNA (the eWLC parameters and experimental data are from Figure 5A of Ref. 8, with the extension values divided by a contour length of 1.15 *μ*m), respectively. Insets: the probability density *ρ* of relative bond lengths *r/r*_0_ at *L*_*x*_*/L*_0_ ≈ 1.4 (panel D), *L*_*x*_*/L*_0_ ≈ 1.325 (panel E), and *L*_*x*_*/L*_0_ ≈ 1.4 (panel F).

In the plateau region of the force-extension curves, the simulations show no formation of distinct (macroscopic) phases in the polymer, as illustrated in Figure 4 (cases 1-2). As shown in Figure 4 (case 3), we also see no phase separation in our model with the adding of long-range electrostatic interactions described by the Debye-Hückel potential.

Our prediction is that long dsDNA and dsRNA molecules do not exhibit (macroscopically) distinct phases in the plateau region at room temperature. We speculate that single-molecule fluorescence resonance energy transfer, with judiciously placed labels, may be used to test the prediction.

## 3 Conclusions

In this work, we have suggested a model that could replace the variants of the WLC in interpreting single-molecule stretching experiment. The model, built on a non-convex potential energy function, is able to reproduce well the force-extension curves of the long DNA and RNA with a single force plateau under low and high tension. The model requires five parameters that can be easily derived from a single force-extension curve using the suggested algorithm. The algorithm is based on an approximation of a force-extension curve by a function describing the minimum-energy state of the polymer chain with its fixed ends.

Although the proposed mesoscopic model has its limitations, including no explicit account for sequence dependence or torsional rigidity, we showed that it (i) accurately reproduces the force-extension curves of long dsDNA and dsRNA with one force plateau, and (ii) predicts that, in the plateau region of the force-extension curves, the polymer does not form distinct phases, regardless of the presence of the long-range potential, providing new insight into the mechanics of the polymers. In the future, it will be interesting to explore time-dependent features of the model, including the possibility of describing hysteresis between stretching and relaxation curves.

## 4 Methods

While both atomistic and coarse-grained approaches can be useful to study the behavior of dsDNA or RNA under tension, we argue that a coarse-grained model is needed to study possible tension-induced phase transitions on long enough time-scales. This is because these effects require investigating chains made of *N* ≫ 1 units; such a unit is chosen as 11 base pairs long. A fully atomistic description would therefore call for fragments of ∼ 1000 base pairs long, which would make long enough simulations extremely challenging at the moment.

Here, all MD simulations are conducted in ESPResSo 3.3.1 package ^56^ with Langevin dynamics at 300 K and an integration time step of 100 fs. Each molecule is modeled as a chain of *N* = 99 beads, Figure 1A, connected by non-linear springs (potential *U*(*r*)). Each bead represents 11 base pairs of DNA or RNA (∼ 1 turn of B-DNA or RNA) and has a diameter of 1 Å and a mass of 7150 Da.

The total potential energy of the chain, *U*_*total*_, has two terms: 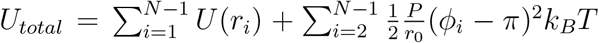. The first term is summed over all adjacent pairs of beads, and *r*_*i*_ is the distance between the centers of bead *i* and bead (*i* + 1). The second term is the harmonic bending energy of the chain (the discrete version of the WLC model), where *P* is its persistence length (in Å); *ϕ*_*i*_, the angle between the vectors from the centers of bead *i* to bead (*i* − 1) and bead (*i* + 1), Figure 1A; the equilibrium bond length *r*_0_ is 11*h*, where the distance between base pairs *h* is given in Table 1 below.

**Table 1:** Simulation details for unconstrained dsDNA (unDNA), torsionally constrained dsDNA (tcDNA), negatively supercoiled dsDNA (spDNA), and dsRNA, where *γ* is the Langevin collision frequency, *K*_*trap*_ is the stiffness of the optical trap and *h* is the distance between base pairs.

| Type | $\gamma^a$<br>(ps<sup>-1</sup>) | $K_{trap}$<br>($k_B T / \text{\AA}^2$) | $h$<br>(\AA) |
| --- | --- | --- | --- |
| unDNA | 0.0254 | 7.3139 | 3.4<sup>b</sup> |
| tcDNA ($\sigma = 0$) | 0.026 | 7.688 | 3.4<sup>b</sup> |
| spDNA ($\sigma = -0.69$) | 0.0163 | 3.027 | 3.7264<sup>c</sup> |
| dsRNA | 0.0198 | 4.457 | 2.9<sup>d</sup> |

<sup>a</sup> We deliberately use much smaller values of Langevin $\gamma$ compared to those of water to dramatically increase the speed of conformational sampling,<sup>57</sup> including the time needed to equilibrate the system. For long DNA fragments, the effective time-scales that can be reached in this low $\gamma$ regime can be up to 100-fold the nominal trajectory length.<sup>57</sup>
<sup>b</sup> B-form DNA.
<sup>c</sup> Equals $3.4k$ , where $k=1.096$ from the eWLC fit for spDNA.
<sup>d</sup> Ref. 8

In Figure 4 (case 3 only), the Debye-Hückel potential, *U*_*DH*_ (*r*) = *l*_*b*_*k*_*B*_*Tq*^2^ exp(−*κr*)*/re*^2^, is added as a perturbation to *U*_*total*_ to model long-range interactions between charged beads in salt solution for *r < r*_*c*_, where *r* is the distance between the centers of beads, *e* is the elementary charge, the Bjerrum length *l*_*b*_ is 10 Å, the charge of each bead *q* is −2|*e*|, the inverse Debye length *κ* is 0.01 Å^−1^, and the force cutoff *r*_*c*_ is 10*r*_0_. At *t* = 0 ns, the contribution of the Debye-Hückel potential to the total potential energy of the chain does not exceed 4 %.

The force in the simulations is measured with a model of the optical trap, represented as a harmonic spring potential, *E*_*trap*_(*r*_1_) = (*K*_*trap*_*/*2)(*r*_1_ − *r*_2_)^2^, where the equilibrium length *r*_2_ is 5*r*_0_, *r*_1_ is the distance between the right fixed end and the center of the rightmost bead of the chain (Figure 1A), and *K*_*trap*_ is the stiffness of the optical trap, Table 1. We make the following assumptions: (i) the average mass of a base pair is 650 Da, (ii) the contour length of dsDNA and dsRNA is fixed at *L*_0_ = (*N* − 1)*r*_0_.

Each simulation is 700 ns long, and begins from the starting conformation of the chain, case 1, 2 or 3 in Figure 4, in which the force in the optical trap is zero, Figure 1A. We find that 700ns is enough to ensure convergence to the same final state regardless of the initial conformation of the chain, Figure 4. To obtain each point on the force-extension curves shown in Figure 3B and 5D-5F, we select the “one-phase” conformation (case 1 in Figure 4) with *r/r*_0_ = 0.8 as the starting conformation of the chain. The duration of each simulation is composed of the 50 ns equilibration run followed by the production run. During the 650 ns production run, the values of the force in the optical trap and the relative extension *L*_*x*_*/L*_0_ are recorded at each integration time step, and then averaged over 650 ns. To obtain the next point on the force-extension curves, the above protocol is repeated starting from the same starting conformation, modified by a 0.025*r*_0_ increment to each bond length. In the insets of Figure 3B and 5D-5F, the relative bond lengths are recorded every 0.1 ns during the 650 ns production run. Table 1 lists other simulation details.

## Acknowledgement

We thank Rodrigo A. Maillard, Graeme A. King for help with interpretation of force-extension curves from optical tweezers experiments, J. Ricardo Arias-Gonzalez for providing raw experimental data points in Figure 5F. We especially thank Igor S. Tolokh for his helpful comments on the manuscript. This work was supported in part by the National Science Foundation (MCB-1715207).

## Graphical TOC Entry

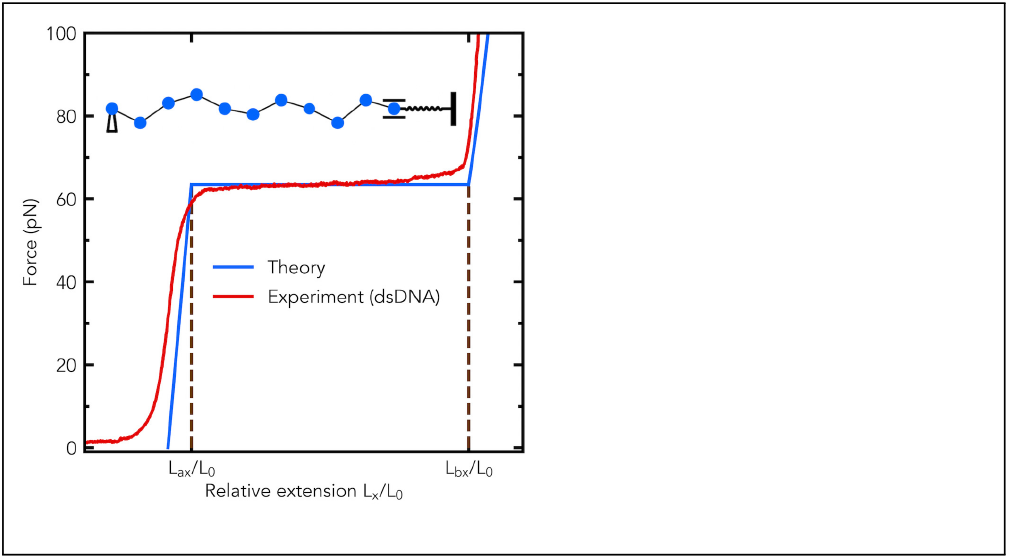

